# Revealing non-trivial information structures in aneural biological tissues via functional connectivity

**DOI:** 10.1101/2024.05.09.593467

**Authors:** Douglas Blackiston, Hannah Dromiack, Caitlin Grasso, Thomas F. Varley, Douglas G. Moore, Krishna Srinivasan, Olaf Sporns, Joshua Bongard, Michael Levin, Sara I. Walker

## Abstract

A central challenge in the progression of a variety of open questions in biology, such as morphogenesis, wound healing, and development, is learning from empirical data how information is integrated to support tissue-level function and behavior. Information-theoretic approaches provide a quantitative framework for extracting patterns from data, but so far have been predominantly applied to neuronal systems at the tissue-level. Here, we demonstrate how time series of Ca^2+^ dynamics can be used to identify the structure and information dynamics of other biological tissues. To this end, we expressed the calcium reporter GCaMP6s in an organoid system of explanted amphibian epidermis derived from the African clawed frog *Xenopus laevis*, and imaged calcium activity pre- and post- a puncture injury, for six replicate organoids. We constructed functional connectivity networks by computing mutual information between cells from time series derived using medical imaging techniques to track intracellular Ca^2+^. We analyzed network properties including degree distribution, spatial embedding, and modular structure. We find organoid networks exhibit more connectivity than null models, with high degree hubs and mesoscale community structure with spatial clustering. Utilizing functional connectivity networks, we show the tissue retains non-random features after injury, displays long range correlations and structure, and non-trivial clustering that is not necessarily spatially dependent. Our results suggest increased integration after injury, possible cellular coordination in response to injury, and some type of generative structure of the anatomy. While we study Ca^2+^ in *Xenopus* epidermal cells, our computational approach and analyses highlight how methods developed to analyze functional connectivity in neuronal tissues can be generalized to any tissue and fluorescent signal type. Our framework therefore provides a bridge between neuroscience and more basal modes of information processing.

**Author summary:** A central challenge in understanding several diverse processes in biology, including morphogenesis, wound healing, and development, is learning from empirical data how information is integrated to support tissue-level function and behavior. Significant progress in understanding information integration has occurred in neuroscience via the use of observable live calcium reporters throughout neural tissues. However, these same techniques have seen limited use in the non-neural tissues of multicellular organisms despite similarities in tissue communication. Here we utilize methods designed for neural tissues and modify them to work on any tissue type, demonstrating how non-neural tissues also contain non-random and potentially meaningful structures to be gleaned from information theoretic approaches. In the case of epidermal tissue derived from developing amphibians, we find non-trivial informational structure over greater spatial and temporal scales than those found in neural tissue. This hints at how more exploration into information structures within these tissue types could provide a deeper understanding into information processing within living systems beyond the nervous system.

## Introduction

Information and its processing are widely accepted to play a central role in biological function [1–4]. This critical role is particularly important in understanding the function of neuronal tissues, and as such, information theoretic approaches have seen wide-spread adoption as quantitative tools in neuroscience [5]. Insights into neuronal function from these approaches include collective decision-making by groups of neurons, how long-range correlations are structured across neural networks, and the structure of phase transitions in networks of neurons to name a few [6–9]. However, communication and information processing are exclusive to populations of neurons: these are embodied processes throughout the cellular architectures of multicellular life [10, 11]. Yet, the application of information theoretic approaches has seen limited development towards a universally implementable approach to quantify general tissue function and behavior to understand the role of information in other tissue types beyond neuronal examples [12–14].

In multicellular systems, cells must collectively coordinate their actions to regulate the diverse range of processes essential to multicellular life: these include regulation of pattern formation in development, morphogenesis, wound healing, regeneration, and behavior among others. Communication and information sharing can even extend beyond species-specific boundaries, as is the case for plant-animal interactions and in symbiotic associations like lichen where multiple species are in direct coordinated communication. Traditional techniques for characterizing coordination in non-neural cellular tissues, thus far, include multielectrode arrays [15, 16], planar cell polarity analysis [17, 18], physiological reporter dyes [19], immunohistochemistry [20], and RNAseq [21, 22]. These have provided important methods for gaining insight into specific cellular function, such as, neural voltages at various cell stages, alignment, and coordination of cells within a tissue plane, distribution and localization of biomarkers, and global ligand and receptor interactions [23]. However, an open challenge is capturing the longer temporal and spatial scales necessary to characterize information processing associated with collective behavior and coordinated decision making across entire tissues and whole multicellular organismal systems.

One approach available to capture these longer-range dynamics is provided by functional connectivity (FC) networks derived from information theoretic analyses of signal data. These networks provide a quantitative framework for identifying connections and information flow over spatial and temporal scales relevant to the coordinated function of entire tissues [24]. FC networks are weighted, undirected networks generated based on instantaneous statistical correlations, or statistical dependencies, between activity in different areas of a system. They are often employed in the field of neuroscience to quantify temporal correlations in activity between different regions of the brain, as in the case of correlated neural firing between regions of the brain which can reveal correlated behavior, even if the regions are spatially separated [25]. Information theory provides a set of mathematical tools to quantify such correlations, where measures such as mutual information [26] and transfer entropy [27] applied across temporally sampled data can be used to construct FC networks that capture nonlinearities and structure not apparent in static images. A predecessor and contemporary approach to constructing FC networks is anatomical connectivity maps, which focuses on physical tracts that can reveal direct anatomical links between different physical regions in a tissue. However, these do not capture the long range temporally correlated structures, which need not be in direct physical contact, which are revealed in FC networks. Therefore, we adopt the approach utilized in the study of networks of neurons to develop applications of the same kind for implementation to other multicellular tissues. We anticipate such studies will provide a complement to existing approaches, by allowing a new window into understanding tissue function through understanding FC over longer spatial and temporal scales.

A commonly used signaling molecule is Ca^2+^, which is found across nearly all living systems. Ca^2+^ is held both inter- and intra-cellularly and can be used for tracking rapid physiological responses to a wide range of events [28, 29]. Tracking Ca^2+^ across a tissue sample therefore provides the opportunity for better understanding information flow and use at the level of cellular networks, including cellular communication and information transfer in the context of basal and perturbed states [30, 31]. Indeed, fluorescent reporters of Ca^2+^ are now widely used within neuroscience to track and quantify cellular function and have formed the foundation of information theoretic analyses to understand information processing in neuronal networks [32, 33]. Ca^2+^ is also known to regulate epithelium healing across diverse model species (fish, chick, frog, mouse, human) [34–37], though most work to date has focused on rapid events (milliseconds to seconds) such as at the time of wounding or neuron firing. Due to this limited observational limit it is not known if long range events exist and/or contribute to the informational structure in a tissue. Herein, we demonstrate evidence of such long-range correlations via FC networks, which suggest in our system either has a storage of memory or long-range coordination of cells in wound healing.

To explore information processing and its relation to function in a non-neural tissue, we expressed mRNA encoding the calcium reporter GCaMP6s in an organoid system of embryonic explanted amphibian epidermis derived from the African clawed frog *Xenopus laevis* [38, 39]. This modified self-assembling system, composed of the developing epidermal cells, was selected for its well characterized cell types and diverse uses in self-organization, cell polarity, stem-cell differentiation, wound healing, human pulmonary disease, and biomaterial science [40–47]. To show an explicit example of tracking whole tissue-level behavior using these approaches, our primary focus is on how cellular networks respond to perturbation by inducing a mechanical puncture wound. Using techniques developed for medical imaging, we stabilize videos of recorded organoids and track intracellular Ca^2+^ over time. From these time series, we construct FC networks using bivariate mutual information between cells to infer internal informational structure in the tissue. Topological and communicational properties of the networks, such as mesoscale community structure, degree distribution, and spatial embedding, were characterized to better understand how epidermal cells respond to perturbation, and what controlling parameters are retained by cellular communities isolated from their host. We find that *Xenopus* ex-vivo tissue self-organizes into non-trivial informational structures that can serve as proxies for the intact organism, displaying a pronounced mesoscale network topology. Furthermore, the organization of the tissue is flexible, restructuring itself in response to the puncture and thereby demonstrating a dynamic response to wounding. While we focus on the application of these methods to epidermal tissue herein, our approach is generalizable to any tissue type and fluorescent signal. In what follows, we outline the process of generating the FC networks, review what structures they reveal, and discuss the future directions for using information theory to uncover larger scale temporal and spatial functional structure in multicellular tissues.

### Constructing functional connectivity networks using experimental data derived from multicellular tissues

To examine the informational structure of a non-neuronal tissue, we tracked calcium transience in a vertebrate model of wound healing, using developing *Xenopus laevis* embryos as our source material. At the Nieuwkoop and Faber stage 2 (4-cell stage, Fig 1A), each of the 4 cells were injected with two mRNA transcripts, one encoding the fluorescent calcium indicator GCaMP6s, the other encoding the intracellular domain of the notch protein (Notch ICD) to inhibit motile-cilia formation on the developing epidermis [48–51]. Knockdown of motile cilia was necessary to prevent rotational movement of the organoid which complicates downstream image registration efforts. After 24 hours of development at 14°C, the animal cap of the embryo was excised manually with surgical forceps (Fig 1B, red circle) and cultured in a saline media. Following an additional 24 hours of development (Fig 1C), the developing spheroid of tissue can be left untreated or compressed (Fig 1D) to produce a flattened morphology amenable to 2D fluorescent microscopy.

**Fig 1.**
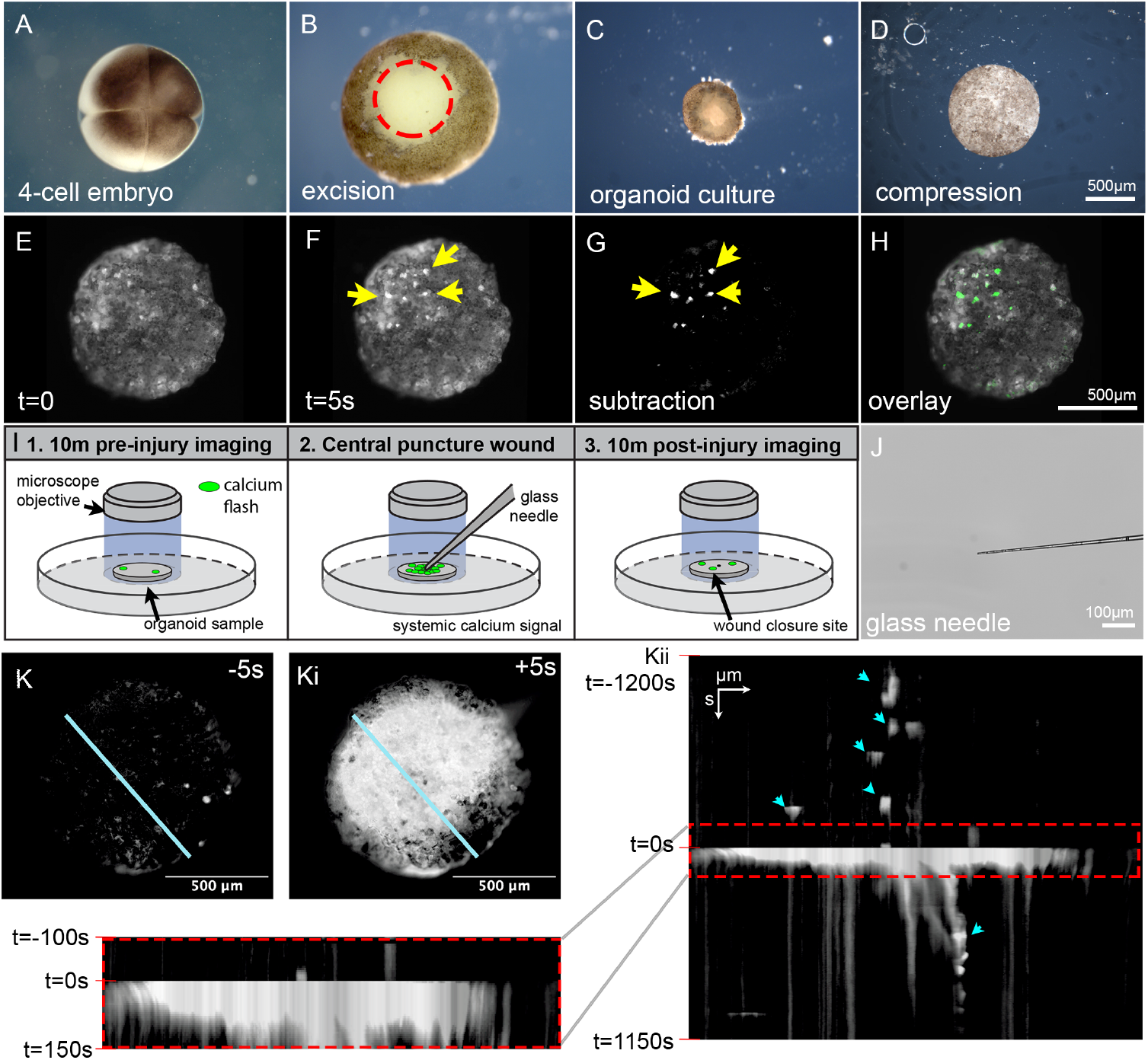
Long-term calcium transience in vertebrate epithelium in the basal state and following injury. Calcium transience within these tissues displays more diverse structures when longer timescales are observed. The kymograph analysis standard timescale is on the order of milliseconds, when applying this to the punctured explant revealed no discernible dynamics even when expanded to the order of hundreds of seconds. When opened to the order of thousands of seconds new structures are observed within the tissue. What these structures represent requires further investigations beyond the scope of kymograph analysis.

Calcium imaging was performed 7 days post-fertilization, at which time the epidermal organoid was fully differentiated, containing 3 distinct cell types on the surface: mucus producing goblet cells, small secretory cells, and ionocytes [40, 41, 52]. Preliminary studies found that a capture rate of 1 frame per 5 seconds was sufficient to identify individual calcium flashes across the surface of the tissue without inducing phototoxicity. Individual frames of the timelapse dataset (Fig 1E) were subtracted from subsequent frames in the stack (Fig 1F) to identify cells presenting calcium flashes (Fig 1G, H). This imaging setup was found to be stable over the duration of 10-20 minutes. The experimental setup consisted of 10 minutes of organoid imaging in its basal state, preceded by a centrally located puncture wound delivered via a pulled glass capillary, followed by an additional 10 minutes of organoid imaging during wound resolution (Fig 1I, J). A total of 6 organoids were imaged in the experimental setup, all at 7 days of development to reduce age related variance in downstream analysis.

Kymographs are frequently utilized to visualize calcium topography following wounding, as time is represented as a dimensional axis. When employed on the organoid injury dataset, systemic calcium activation is readily observed, and resolved, over the course of 100s (Fig 1Ki, ii, red box). The sharp transition noted at 0s (Fig 1Kii) is an artifact of imaging, as the time series omitted the moment of puncture when the needle occluded the optics, and re-centering the sample was necessary following injury. This method proved sufficient to capture the large-scale calcium changes in direct response to injury, matching previous reports of lacerations in the same system [53]. Interestingly, when kymographs were expanded beyond the standard time frame convention prior and post injury by the order of thousands of seconds, different structures could be observed in the data in the form of less frequent flashes by individual cells (Fig 1Kii, teal arrows). This is the first evidence of non-trivial long-range correlations within the informational structures of non-neural tissues.

Statistical analysis of these flashes to determine structure is limited in kymograph representations, due to the nature of the linear slices used in the method (Fig 1Ki, teal line) which occluded the less frequent signals. Thus, to analyze this data for more complex information structures prior to, and post injury, FC maps were utilized for whole-image analysis. We therefore performed a coarse-grained analysis and visual inspection of the global calcium signal, which revealed a sharp increase in signal at the time of puncture (*t* = 0s in Fig 2A, B). The signal remained high and unstable for a period following puncture, which varied in duration across organoid samples. During this time, organoids shifted in position due to force imparted from the glass capillary. These movements were too great to correct for using conventional image registration software and were thus excised from the video. Resulting in two distinct videos per organoid: one capturing the basal state behavior prior to puncture damage (*pre-puncture*) and the other capturing behavior post damage once the organoid had settled (*post-puncture*). Pre- and post-puncture videos were processed and analyzed independently. Smaller translational and rotational movements between frames were corrected using ANTs image registration software in *Julia* [54]. After organoid alignment, temporal averaging of the images was used to produce a representative image that could be supplied to the deep learning cell segmentation model, Cellpose (Fig 2C, G) [55]. Preliminary experiments explored optimal model parameters for each organoid, however, performance was highly dependent on image quality. Models performed suboptimal in regions that were out of focus and/or of high fluorescent intensity where cell boundaries were obscured. Thus, the segmented cells identified were spread non-uniformly across the organoid. Pixel intensities within each identified cell boundary were extracted and averaged in each frame to produce a time series of calcium readings localized to individual cells at every time step (Fig 2D, H).

**Fig 2.**
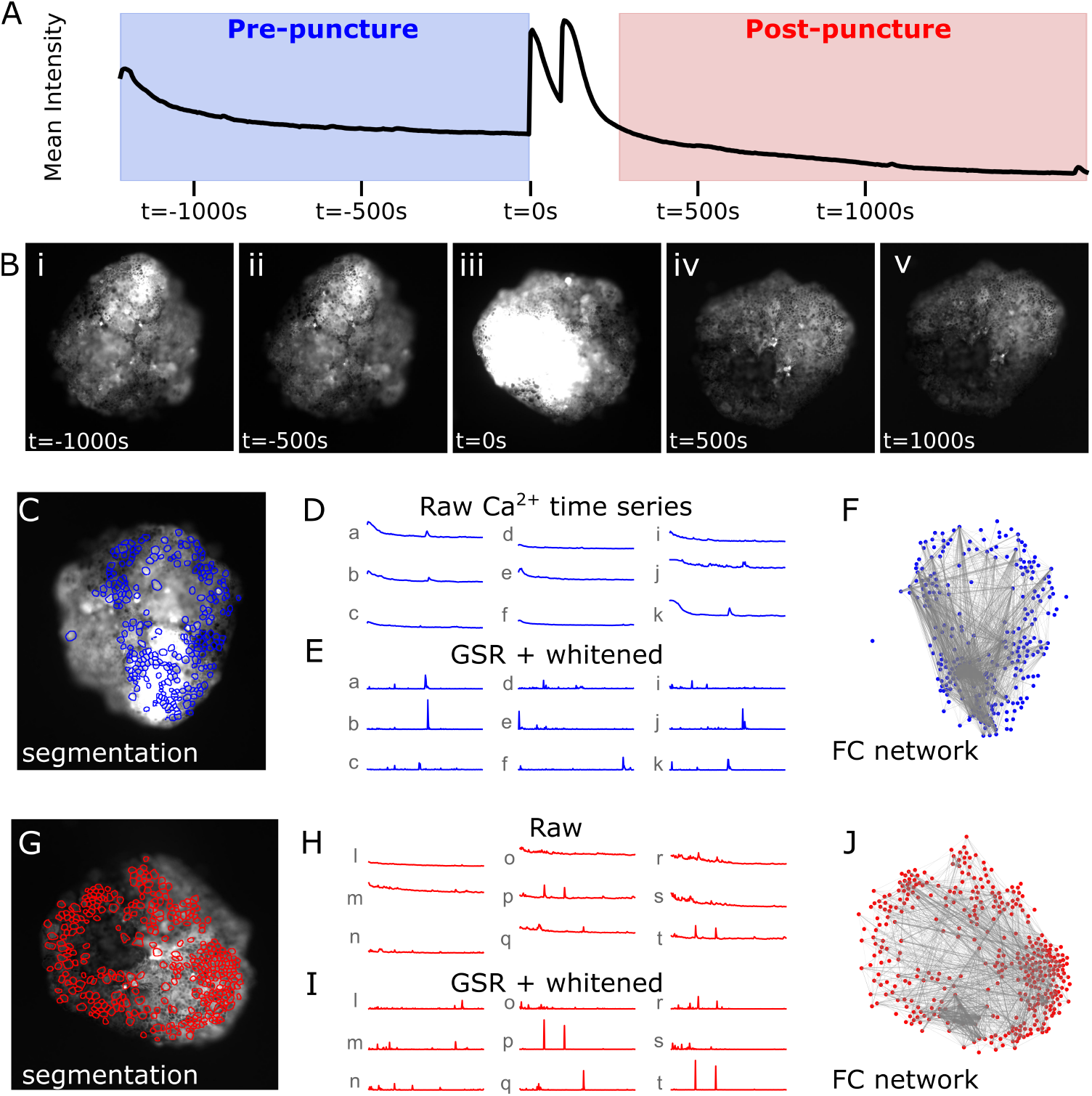
FC inference pipeline as observed in organoid 4. A: Average pixel intensity over time, observed signal peaks at the time of puncture (t = 0s) and remains unstable for a period post puncture. Frames at the time of puncture are removed, producing two distinct videos: pre-puncture (blue) and post-puncture (red). B: Calcium transience throughout the observation time frame. Orientation of the organoid changes due to impact from the needle at t = 0s. C: Cell segmentation as determined by Cellpose pre-puncture. D: Raw calcium signal intensity time series for a random sample of nine segmented cells (a-k) pre-puncture. E: Global signal regressed (GSR) and whitened calcium intensity time series for the same cells (a-k). F: FC networks are generated by computing mutual information between all pairs of cells’ GSR + whitened calcium signal intensity time series. Nodes of the network (blue dots) represent segmented cells in the organoid. Edges of the network (gray lines) represent non-zero mutual information between a given pair of nodes. G-J: Same as C-F but for the post-puncture video and displayed in red.

Some common problems in analyses such as these arise in managing global artifacts and autocorrelation. Global artifacts, observed in many organoids, include the steady decline in signal at the beginning of pre- and post-puncture videos. To remove this, global signal regression (GSR) is used; the global signal, acquired by taking the average of all cells’ time series, is subtracted from each cell’s individual time series. Additionally, to remove any autocorrelation, time series were “whitened” by computing the local conditional entropy rate, a measure of instantaneous information that cannot be learned from observing a cell’s own past signals [56, 57]. These preprocessing steps produce flattened time series with emphasized flashes where there are changes in signal not resulting from autocorrelation or a global artifact (Fig 2E, I). Here, FC is computed as the temporal correlation in activity between pairs of identified cells where correlation is measured as non-zero significant mutual information and activity refers to calcium intensities localized to individual cells. This translates to how much information the flashing pattern of one cell discloses about that of other cells; high functional connection indicates that observing the calcium signal of one cell in a pair provides a lot of information regarding the signal of the other. FC networks thus represent the intrinsic signaling dynamics of a given system over the entire spatial and temporal scale available for analysis. FC network architecture of the epidermis tissue was examined at both the basal state and in response to an injury. Nodes represent identified cells in the organoids and edges represent the magnitude of functional connection (Fig 2F, J). Investigating their properties and organization yields insights into the information structure of this complex, non-neural tissue, and the differences in structure pre- and post-puncture perturbation.

### Functional connectivity networks pre- and post-puncture

FC is a time-averaged, pairwise measure of correlation. Unraveling this measure in the time dimension produces an *edge time series* of instantaneous correlations between pairs of cells’ signals. Edge time series of post-puncture networks were seen to have more highly correlated edges at the beginning of the post-puncture observation period (Fig 3, top row of quadruplet plots). This is supported by the root sum square (RSS) amplitudes which represent the combined magnitudes across all edges for a given network (Fig 3, bottom row of quadruplet plots). Pre-puncture RSS amplitudes do not display a consistent trend across organoids. Post-puncture RSS amplitudes (except for organoid 2) display a consistent trend in which amplitude is high at the beginning of the observation window before rapidly declining towards a baseline. Taken together, these results suggest increased integration among cells via highly correlated Ca^2+^ signals soon after undergoing puncture damage with quick stabilizing back to a baseline. Interestingly, some of the edge-time series display bands of high-amplitude, global co-fluctuations (Fig 3, O2-post, O4-pre, O6-pre), known as *events* in neuroscience. These intermittent episodes have previously been observed in human brain data [58] and are linked to the presence of a complex underlying generative structure in the anatomy [59]. The significance of seeing high amplitude *events* in organoid tissue is currently unclear, but we propose that it may be a fingerprint of a non-trivial interaction structure among the cells as the anatomy of the tissue is nearly uniform and organized similar to a checkerboard, with alternating cell types at regular intervals. Further research is planned in this area.

**Fig 3.**
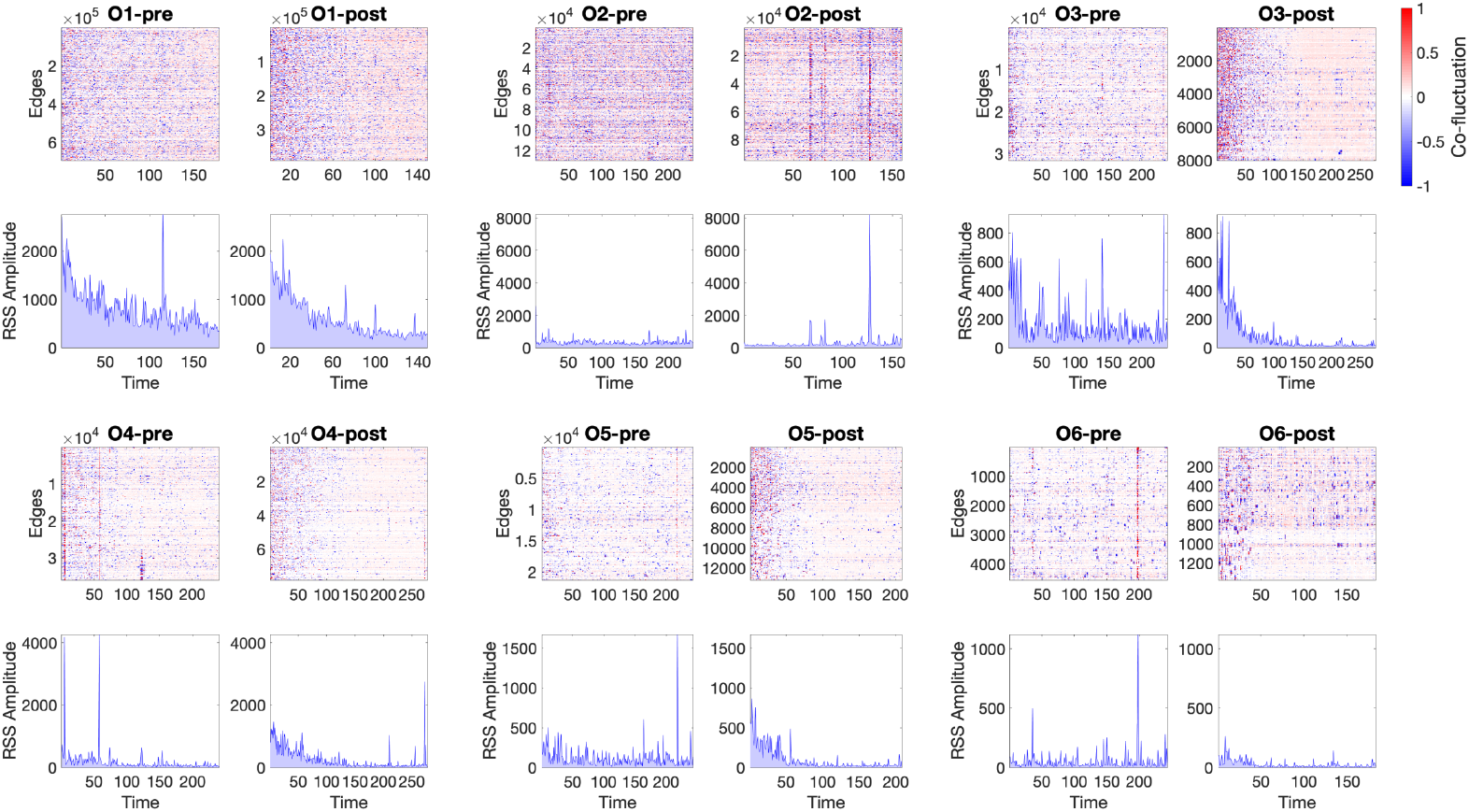
Edge time series, computed as the element-wise product of two z-scored calcium time series, measures instantaneous co-fluctuation between pairs of nodes pre- (left) and post- (right) damage for each organoid (top row in quadruplet). Co-fluctuations are plotted by the magnitude away from the mean; red signifies is above the mean, blue is below, and are interpreted as how strongly the cells are functionally connected. Root sum squared (RSS) amplitude shows the points in time where many cells collectively co-fluctuate (bottom row in quadruplet). In pre-puncture networks there is no clear pattern in co-fluctuations across the organoids, though in post-puncture networks there is a general decrease from strong co-fluctuations and amplitudes to some baseline levels.

Network analyses were implemented to interrogate and characterize the structure of the FC networks, including computing network density and number of edges and nodes (S1 Fig). Degree distributions give the number of edges connected to a given node providing a good corollary to global network structure. Networks both pre- and post-puncture display degree distributions with heavier tails and higher maximum degree as compared to our null model (Fig 4). Null models were constructed for each network by averaging degree distributions from an ensemble of 100 Erdő R*è*nyi random graphs, constructed with the same number of nodes and edges as the corresponding empirical network. These differences between the empirical and null models suggest that networks both pre- and post-puncture are not random, but rather contain nodes that are much more connected than expected by random chance (i.e., the empirical network contains hub nodes). Correspondingly, there are also more nodes with fewer connections than expected by random assignment of edges. Furthermore, Kolmogorov-Smirnov tests revealed pre- and post-puncture distributions were significantly different from one another, except for organoid 6, which contained significantly fewer nodes post-puncture than the other networks (Table 1, S1 Fig).

**Table 1.** Kolmogorov-Smirnov (KS) test statistics and Bonferroni corrected p-values between pre- and post-puncture distributions.

|  | KS | *p* |
| --- | --- | --- |
| O1 | 0.095 | 0.001 |
| O2 | 0.367 | 0.000 |
| O3 | 0.534 | 0.000 |
| O4 | 0.243 | 0.000 |
| O5 | 0.270 | 0.000 |
| O6 | 0.188 | 0.928 |

**Fig 4.**
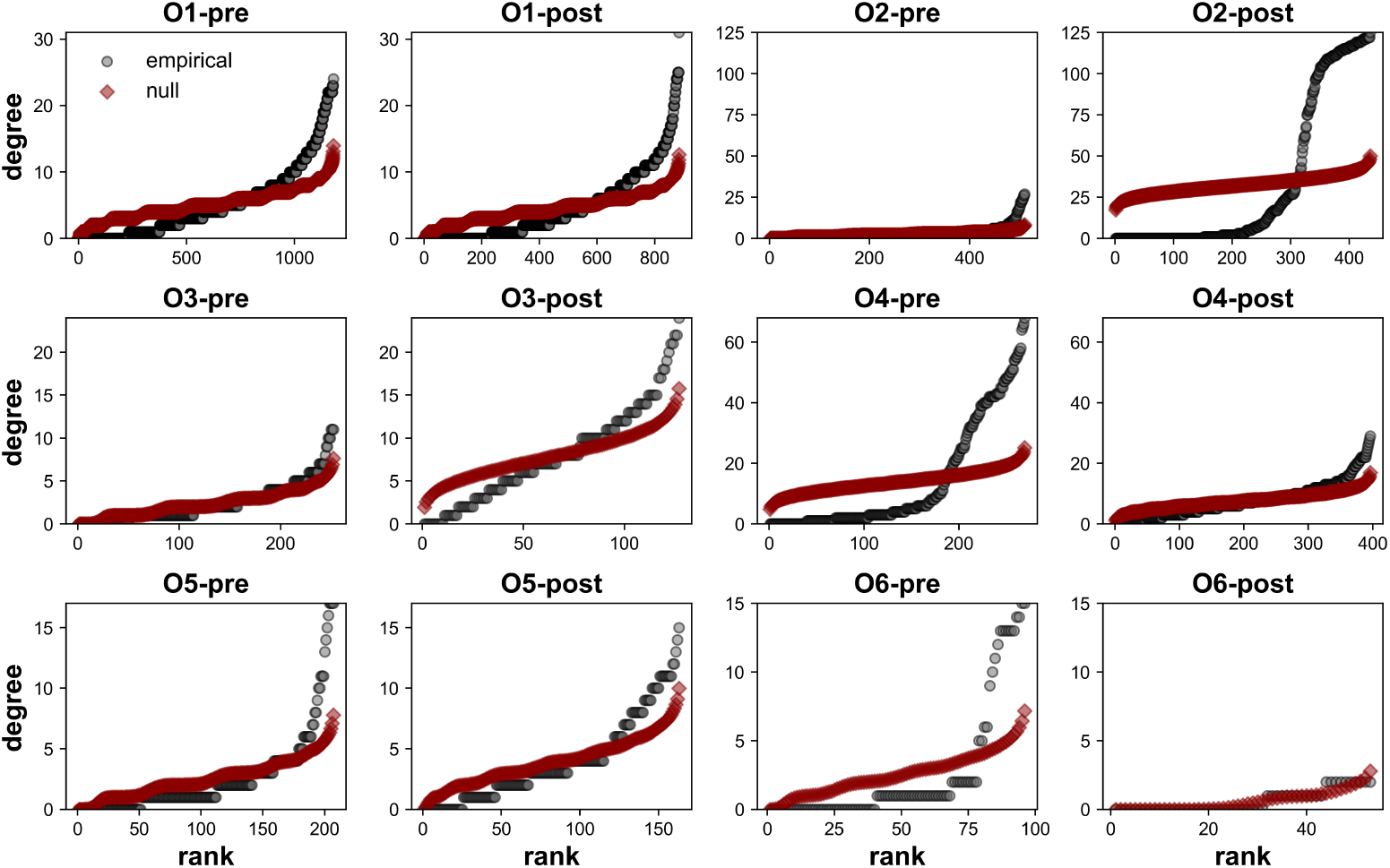
Degree distributions. Empirical networks (black) have heavier tails and higher maximum degrees than expected by random networks (red). The null network model used is the average of 100 Erdo őRènyi graphs with the same number of nodes and edges as the corresponding empirically derived network.

To illuminate the underpinning characteristics of the non-random structure of these networks, the correlation between the magnitude of functional connection and spatial distance between node pairs was investigated via Spearman correlation (Fig 5A). We found significant negative correlations between the bivariate mutual information between pairs of cells and the distance separating those two cells across organoids pre- and post-puncture, except for post-puncture networks for organoids O5 and O6 (negative Spearman correlation coefficients in Fig 5B, S1 Table). That is, our data supports a conclusion that spatially closer cells generally have more coordinated signaling patterns. Such a finding is in accordance with known signaling in non-excitable tissues, in which adjacent cells are connected structurally via an extracellular matrix and extracellular ligand-receptor interactions, and internally via gap junctions allowing the passage of small molecules between neighbors [60, 61]. Spearman correlation coefficients post-puncture appear less negative than their pre-puncture counterparts suggesting an increase in higher magnitude long range connections; however, *N* = 6 is likely too small a sample size to extrapolate to general trends.

**Fig 5.**
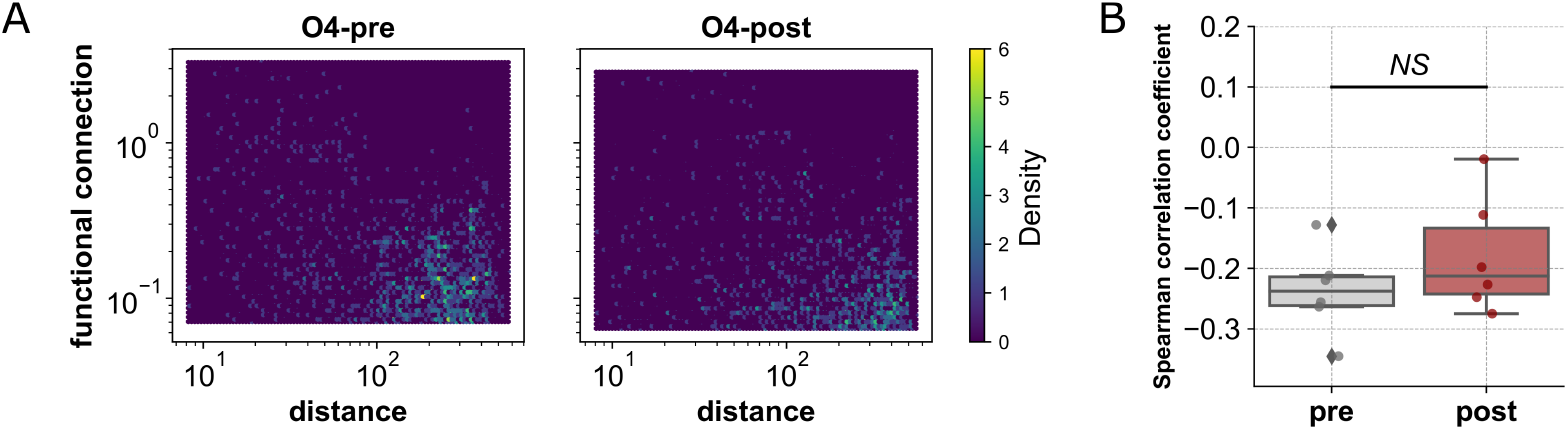
FC is negatively correlated with distance between nodes. A: Spatially closer nodes in organoid 4 tend to have a higher FC, represented by higher density on the plot in yellow, pre- (left) and post- (right) puncture. Organoid 4 has a Spearman correlation coefficient of −0.22 (*p* = 0.00) before damage and −0.18 (*p* = 0.00) after damage. B: Spearman correlation coefficients pre- and post-puncture for all *N* = 6 organoids.

Another method we implemented to uncover non-trivial structure is community detection, which reveals the modular nature of the networks by identifying groups of cells that are highly functionally connected regardless of physical location. Community detection was performed using multiresolution consensus clustering [62] with the Louvain method [63, 64]. Such clustering algorithms work by organizing nodes into groups that maximize the number of within-group edges and minimize the number of between-group edges. FC matrices, square matrices with nodes on both axes (*i, j* ∈*V*) and entries colored by the magnitude of functional connection between nodes *i* and *j*, are sorted by modular structure, placing nodes within the same community next to one another on the axes. Thus, modules differentiate as squares along the diagonal with high levels of FC (Fig 6A). Modules can be interpreted as clusters of cells with large statistical dependencies between cells within the cluster compared to those outside the cluster. The cause of such integrated clusters is not readily obvious; however, as shown in Fig 5, spatially closer cells appear to have more correlated signaling dynamics and thus modules might appear as groups of cells clustered in space. Unexpectedly, this is not always the case. Spatial visualization of the three largest communities both in the organoid (Fig 6B) and the FC network (Fig 6C) reveal that while some communities are indeed clustered in space (Fig 6B, C, green and orange modules), others contain nodes that are spread across the entire organoid (Fig 6B, C, blue module). This is further emphasized by observing the distribution of within-module and between-module distances, normalized by the maximum distance between two cells in the organoid, across all samples (Fig 6D). Within-module distributions both pre- and post-puncture have clear peaks at small distances but also extend to larger distances – a noticeably distinct shape compared to the smooth normal distribution formed by between-module distances. This unique shape both validates the existence of spatial clustering of communities across organoids but also suggests modular structure is not entirely spatially dependent and there may be non-trivial functional correlations present in the more spread-out modules. To further uncover underlying characteristics of this modular structure, we employed a participation coefficient measure in which each node is scored based on the number of unique communities its neighbors in the FC network belong to [65]. Nodes in post-puncture networks are connected to a significantly higher fraction of modules (i.e. have more diverse neighbors) than before puncture, again hinting at an increase in integration among cells following puncture perturbation (Fig 6E).

**Fig 6.**
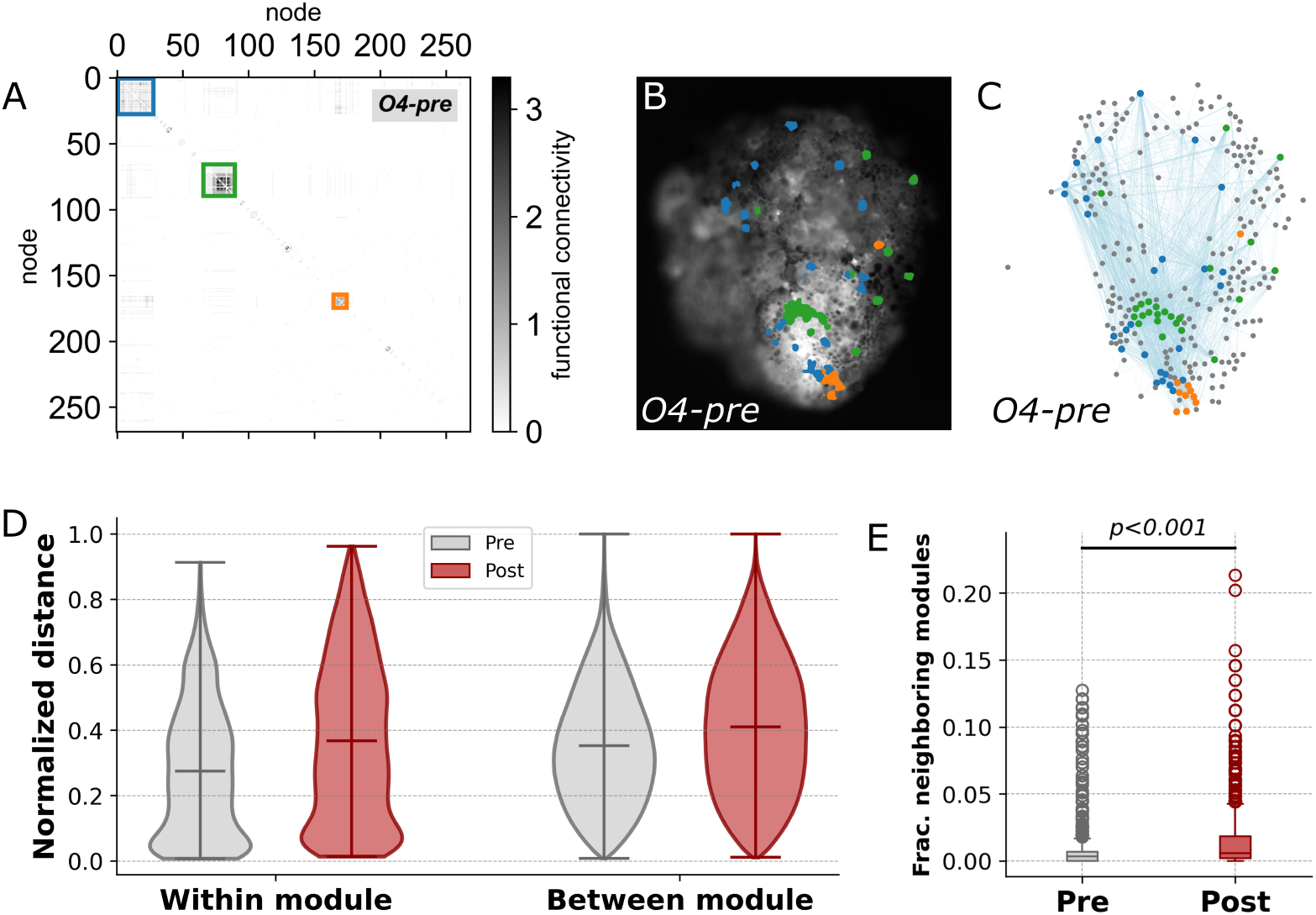
Network modularity. A: FC matrix reordered by modular structure for organoid 4 pre-puncture with the three largest communities highlighted (blue, green, orange, largest to smallest). B: Physical location of nodes composing the three largest modules painted by color in organoid 4 pre-puncture at *t* = −1220s. Modular structure can be clustered locally in space, distributed across the spatial extent of the organoid, or have a combination of the two. C: FC network visualization with nodes placed by physical location and painted by corresponding module color (gray represents all other nodes not in the three largest modules). Nodes within a module have more connections to other nodes within the same module than to nodes outside the module. D: Within- and between-module distance distributions pre- and post-puncture, normalized by the size of the network for *N* = 6 organoids. E: Neighborhood modular diversity, measured as the number of distinct modules a given node’s neighbors are members of, post-puncture networks showed significantly higher diversity then pre-puncture networks (Mann-Whitney U test, *U* = 2004296, *N* = 6, *p <* 0.001).

## Discussion

Here we have demonstrated the use of a general, information-theoretic approach for the large-scale spatiotemporal analysis of functional connectivity in a non-neural organoid system. Mutual information was employed as a measure of the correlations between segmented cells’ calcium transience in an organoid as it experiences, and subsequently recovers from, a puncture wound. The approach utilizes information-theoretic and regression preprocessing techniques to control for autocorrelation and global artifacts in the data, ensuring meaningful correlation is captured in the resulting FC networks. We find non-trivial, non-random FC network structure consistent across organoids both pre- and post-puncture. These networks possess characteristics well-known in other biological systems such as heavy-tail degree distributions and banding behavior, corresponding to *events* of high amplitude co-fluctuations.

Consistent with what is known about calcium signaling in epidermal tissue, we find that the functional connection between cells shows negative correlation with the distance between them. This, however, does not preclude the presence of long-range, high-magnitude functional connections. In fact, modular analysis of the networks revealed the existence of highly correlated, spatially diverse communities of cells. The cause of such modules could be explained by several features, including the three cell types spaced at regular intervals across the organoid surface, which may have individual calcium dynamics, or the propagation of non-observable signals below the outer layer of cells. However, discriminating between these possibilities will require additional analyses that will be the topic of future work.

In the face of perturbation, networks retain key characteristics defining their structure such as heavy tail degree distributions and the presence of hub nodes, as well as similar spatial embeddings and modular structures pre- and post-puncture.

Interestingly, however, our results suggest an increase in integration among cells shortly post-puncture as evidenced by the heightened correlation at the beginning of the post-puncture edge time series. Such a finding suggests that the information structure is dynamic and able to restructure itself in response to damage. However, we were unable to discern any reliable signatures of this phenomenon from the data. For example, these might have included statistically significant results indicating an increase in high-magnitude long-range connections post-puncture. Or subtle differences in modular structure, which we could not identify due to limitations in this study, such as the small sample size of *N* = 6 organoids and relatively short time series.

An additional limitation in the approach we present is the requirement of the segmentation of individual cells over time from fluorescent microscopy data – a notoriously challenging task. Image segmentation algorithms are susceptible to poor performance when cell boundaries are obscured due to blurry or out of focus regions of the image or regions in which there is widespread high intensity (i.e., a tissue wide calcium flash). The recordings used in this study suffer from a combination of these two challenges. Furthermore, the mechanical puncture event displaced the organoids, preventing image registration software from being applied to an entire event by recording it continuously. Thus, preprocessing and analysis of pre- and post-puncture observation periods of the organoid were carried out in isolation leading to pre- and post-puncture videos having inconsistent segmented cells. Comparing networks with different numbers and placements of nodes renders direct comparisons between pre- and post-puncture networks challenging and thus, we were severely limited in the analyses we were able to perform. A focus of future work will be on experimental methods for less disruptive perturbations to enable tracking of the sample throughout the entire observation period. Despite these limitations, we have displayed the potential of these tools and believe that with more data at our disposal this could be a very powerful and comprehensive approach to non-neural tissues.

Aside from increasing the quality and quantity of data, future work will explore the use of other measures of dependency between cell activity: the FC approach is undirected and does not account for time-directed effective connections (where the past state of one cell influences the future state of another). Measures of effective connectivity such as the transfer entropy may provide a more refined perspective on information “flow” by considering temporal directionality of signals [27]. Furthermore, there has recently been an explosion of interest in the phenomena of higher-order/beyond-pairwise interactions in complex systems [66, 67], and many of the tools that have been developed could be easily slotted into the general framework we present here [68–70]. We outline a flexible approach to the problem of inferring structure from data, and prospective users have considerable freedom to tailor the approach to different notions of “structure”, including directed or undirected, temporal or atemporal, pairwise or higher-order, and so-on.

Biologically, the use of information theory can identify long-term signaling dynamics within tissues to serve as the basis for developing and testing hypotheses about the nature of information processing and its relation to whole-tissue function. While no mechanistic biological claims are made from the data presented herein, significant changes in network modularity, including neighborhood diversity, can be observed pre- and post-injury. Are these changes instructive or merely an epiphenomenon of the healing process? Suppression, or enhancement, of these networks via calcium signaling activators and inhibitors could help shed light on this question [71]. Other related questions include: how do network dynamics change in the face of different types of injury, from mechanical, to thermal, to chemical? What are the relative contributions of each cell type to the network dynamics? Are correlated longer-range *events* associated with specific cell types? Do mutations mimicking human disease alter network modularity in addition to tissue function? Are network dynamics evolutionarily conserved among diverse taxa, or do they diverge between related species? Or across individuals within a species? All of these are important questions worth exploring, though each may not require the generation of novel tools and methods. The described approach, along with a growing palette of complementary computational tools, shows an avenue that currently available tools are generalizable to diverse biological systems and will likely reveal several hidden signaling modalities across them currently unexplored or understood.

## Materials and methods

### Animal Husbandry

All experiments were conducted using tissue sourced from the amphibian *Xenopus* laevis. Wild type embryos were collected 30 minutes post-fertilization and raised in 0.1x Marc’s Modified Ringer’s solution (MMR), pH 7.8, until microinjection at the 4-cell stage and animal cap excision at Nieuwkoop and Faber stage 9 [72]. Experimental procedures using animals for experimental purposes were approved by the Institutional Animal Care and Use Committee and Tufts University Department of Laboratory Animal Medicine under protocol number M2020-35.

### Microinjection

Microinjection of synthetic mRNA was performed at the 4-cell stage using a pulled glass capillary, with each of the 4 cells being injected to ensure ubiquitous expression across the embryo. Synthetic mRNA was synthesized from a linear DNA template using commercially available kits (Life Technologies), which was stored at −80°C until used. Directly prior to injection, cohorts of healthy wild type embryos were transferred to a laser etched petri dish containing 3% Ficoll solution. The 4 individual cells of each embryo were then injected with a pulled glass capillary, delivering approximately 500 ng of mRNA in 50nL of volume to each cell. After healing for 1 hour, the embryos were washed twice in 0.1x MMR, pH 7.8, to remove the Ficoll solution, and any damaged embryos were discarded before moving the dish to a 14°C incubator. Two mRNA’s were co-injected in the reported work; GCaMP6s, a reporter of calcium activity [73, 74], and the intracellular domain of Notch (Notch ICD), which is known to inhibit multiciliated cell induction in developing frog epidermis [50, 75, 76]. Multiciliated cells were molecularly inhibited in the current study as the presence of these motile structures causes the mucociliary organoid to move during observation, complicating image analysis [77–79].

### Modified organoid generation

At Nieuwkoop and Faber stage 9, the animal cap of each embryo was removed to generate epidermal organoids. Cohorts of injected embryos were transferred to a Petri dish containing 0.75x MMR, lined with 1% agarose to reduce cell/tissue adherence. Using a pair of sharpened microsurgery forceps, the vitelline membrane of each embryo was removed, and the animal cap (the central portion of the pigmented top of each embryo) was surgically excised and inverted in the dish. These explants are known to develop into irregular epidermis if untreated [38–40, 78, 80]. Following excision, the remainder of the embryos were discarded, and the tissue was allowed to heal into a spheroid over the course of 3 hours at room temperature. Following healing, the developing tissue moved to new dishes containing 0.75x MMR and 5 ng/*μ*l gentamicin, lined with 1% agarose, and placed back at 14°C. After an additional 24 hours of development, the animal caps were placed under a glass coverslip for 3 hours at room temperature, generating continuous compression, which resulted in a permanent flattened tissue which improved optical measurements. Following compression, the explants were kept at 14°C for 5-6 further days of development until imaging, at which point the tissue had differentiated into a modified epithelial organoid.

### Imaging

All calcium imaging was performed on an Olympus BX-61 microscope equipped with a Photometrics CoolSNAP DYNO CCD camera and CoolLED pE-300 light source. Individual organoids were placed in a depression slide containing 0.75x MMR under a 4x objective. Images were captured using a FITC filter at a rate of 1 frame every 5 seconds, across a total 20 minutes of observation. Capture rate was determined by pilot studies which identified the minimum time scale to record calcium flashes in individual cells, while also minimizing exposure to illumination to avoid photobleaching and/or phototoxicity. For the first 10 minutes, basal rates of calcium activity were recorded. After 10 minutes the image capture was paused, and a pulled glass needle with a tapered tip diameter of 10-15*μ*m was used to place a puncture near the center of the organoid. Tip diameter was chosen to minimize overall damage to the organoid, and the depth of the wound traversed the entire width of the tissue. Immediately following injury, image capture was reinitiated, and proceeded for an additional 10 minutes of observation. Each organoid was imaged, and injured, individually before being transferred to a new dish, separate from the samples awaiting processing. Between each observation period, the glass depression slide was washed with distilled water, cleaned with a Kimwipe (Kimtech Science), and loaded with fresh 0.75x MMR to avoid sample contamination across trials. Organoids were imaged across two successive days of development, corresponding to Nieuwkoop and Faber embryonic stages 37-40. All images were captured in tiff format and combined into AVI video files for computational analysis using the FIJI software package [81].

### Video preprocessing

A series of preprocessing steps were performed to transform the videos into time series of calcium intensities for each identified cell in the organoids over time. The puncture event caused extreme movement of the organoids as well as a high intensity flash of calcium across the entire organoid obscuring cellular boundaries, making it difficult to track cells throughout the course of the entire video. Full videos were therefore separated into two distinct videos: pre- and post-puncture event. The end of the pre-puncture video was aligned with the time of puncture and the start of the post-puncture video was aligned with the end of immediate high intensity flash that the puncture caused. Image registration was performed on each pair of videos to correct for rotational movement of the organoids, improve the quality of video with a flatfield correction, and do any necessary video cropping. This process was carried out using Advanced Normalization Tools (ANTs) software [54]. Motion correction aligns cells in the organoid throughout time such that a segmentation algorithm can be applied to the time-average of all frames in the video to obtain cell boundaries for the entire series. Cellpose [55], a generalized deep learning model, was used for cellular segmentation. Hyperparameters of the Cellpose model were tuned for each video based on visual inspection (cell diameter = 15, cell threshold = −2.0, flow threshold = 0.8, resample = False). Pixel intensities within each cell boundary were extracted and averaged at each frame to produce a time series of calcium intensities. These steps result in two time series arrays (pre- and post-puncture) of size # cells× # frames for each organoid (S1 Fig, S2 Table).

### Signal preprocessing

Utilized here was an information-theoretic pipeline to infer pairwise statistical dependencies between individual cells. Information theory has been previously discussed as a general framework for inferring systems-level structures in complex, biological systems [30, 82]. Two signal preprocessing steps were applied to emphasize underlying structures in the data and allow for more meaningful inferences: global signal regression and data whitening. Global signal regression attempts to remove global artifacts by regressing out the mean signal across all cells. Data whitening involves removing autocorrelation from the time series. Calcium imaging data is highly autocorrelated [83, 84] which can reduce the effective degrees of freedom between interacting variables, complicating the inference of dependencies between signals [85]. To address this, we took a whitening approach inspired by Daube *et. al*., 2022 [56]: the calcium data for every cell was transformed into time series of the instantaneous local entropy rate, which contains only that information that is not disclosed by the cell’s own past.

For a given cell, *X*, at every time *t*, the information content of the observation *x*_*t*_ is given by the local entropy:

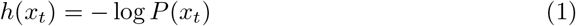

Where *P* (*x*_*t*_) is the probability of observing *X* = *x*. The local entropy (also called the Shannon information content, or *surprisal*) quantifies how much information about the state of *X* is disclosed by the observation of *x*_*t*_. This information can be decomposed into two components:

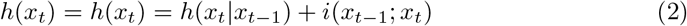

where *i*(*x*_*t−*1_; *x*_*t*_) is the information about *x*_*t*_ that could be learned by observing the immediate past *x*_*t−*1_ (sometimes called the local active information storage [86, 87]), while *h*(*x*_*t*_ |*x*_*t−*1_) is the remaining information that could not be learned by observing the past (sometimes called the local conditional entropy rate [86]).

By transforming the calcium imaging time series into a series of local conditional entropy rates, we effectively *whiten* the signal: every moment is independent of the moment prior to it and contains only that information that could not have been learned by observing *X*’s own past. This remaining information could come from two places: intrinsic randomness in *X*’s own dynamics, or from perturbation by another cell *Y*, whose activity informs on *X*’s own activity. All local entropies were estimated using Gaussian estimators and computed using the JIDT [88] and IDTxl [89] packages.

Finally, after whitening, excessively noisy frames associated with recording artifacts were deleted. Frames where the absolute value of the change in local entropy rate were greater than two times that standard deviation were classified as outliers and removed. The classifications were manually checked by visual inspection as well, to ensure only artifact frames were removed.

### Functional connectivity network inference

Undirected networks for each time series were generated based on instantaneous correlation (functional connectivity) [25]. Nodes of these networks are identified cells and edges are functional connections between each pair of cells in the network computed as the Gaussian mutual information between the pair’s signal time series.

Gaussian mutual information was computed based on the identity:

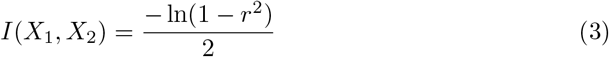

where r is the Pearson correlation coefficient between *X*_1_ and *X*_2_. The transformation into mutual information was chosen as the mutual information, unlike the Pearson correlation coefficient, is strictly non-negative, a key desiderata when analyzing networks. Edges were retained only if the significance associated with the mutual information computation was greater than or equal to *α* ≤ 10^*−*3^ (Bonferroni-corrected against the number of possible edges in the network).

### Co-fluctuation & edge time series

The co-fluctuation analysis was done following the method described in Zamani Esfahlani *et. al*., 2020 [58]. Briefly, each pair of nodal time series was z-scored and multiplied together elementwise to construct an edge time series, where the value of the series at a given time reflects the degree to which those two edges were co-fluctuating together or in opposite directions. Then, the root sum squared deviation from the mean was computed framewise to identify how global co-fluctuations are distributed throughout the duration of the recording (see [59], for more details on high-amplitude co-fluctuations).

## Supporting information

**S1 Fig.**
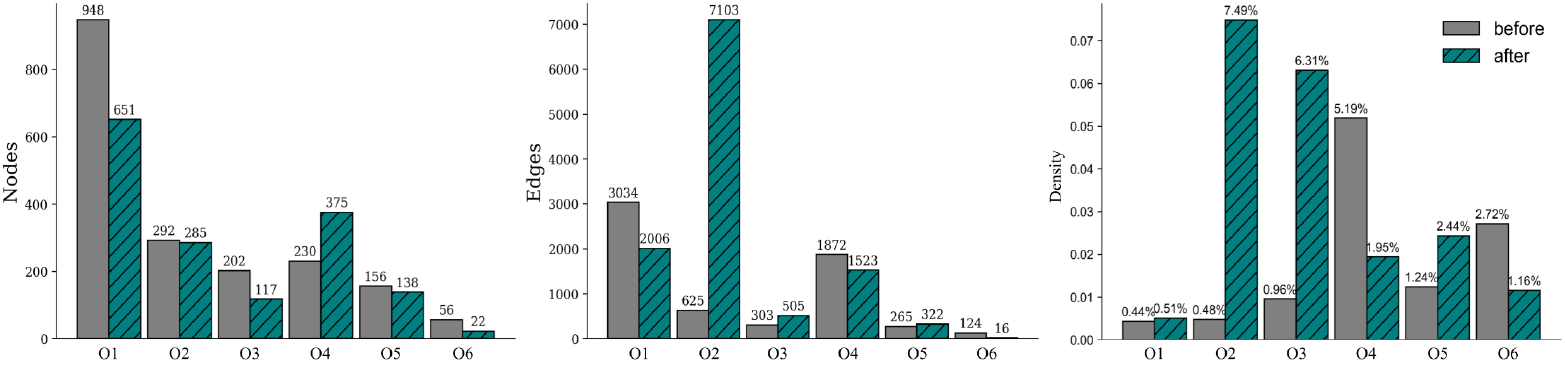
General Network Characteristic.

**S1 Table.** Spearman’s rank correlation between FC and distance between nodes.

|  | pre- puncture |  | post- puncture |  |
| --- | --- | --- | --- | --- |
|  | R | p | R | p |
| O1 | -0.21 | 0.000 | -0.20 | 0.000 |
| O2 | -0.35 | 0.000 | -0.14 | 0.000 |
| O3 | -0.13 | 0.026 | -0.18 | 0.000 |
| O4 | -0.22 | 0.000 | -0.18 | 0.000 |
| O5 | -0.26 | 0.000 | **0.01** | **0.793** |
| O6 | -0.26 | 0.003 | **-0.19** | **0.471** |

**S2 Table.** Number of frames per video. Corresponds to number of time steps in time series.

|  | Pre- | Post- |
| --- | --- | --- |
| O1 | 193 | 151 |
| O2 | 240 | 161 |
| O3 | 241 | 282 |
| O4 | 245 | 279 |
| O5 | 246 | 210 |
| O6 | 246 | 185 |

## Acknowledgments

This research was sponsored by the Defense Advanced Research Projects Agency (DARPA) under Cooperative Agreement Number HR0011-180200022, the Lifelong Learning Machines program from DARPA/MTO. The content of the information does not necessarily reflect the position or policy of the government, and no official endorsement should be inferred. Approved for public release; distribution is unlimited. This research was also supported by the Allen Discovery Program through The Paul G. Allen Frontiers Group (12171) and the Alfred P. Sloan Foundation Matter-to-Life program (G-2021-16495). Additionally, this material is based upon work supported by the National Science Foundation Graduate Research Fellowship Program under Grant No. 1842491. Any opinions, findings, and conclusions or recommendations expressed in this material are those of the author(s) and do not necessarily reflect the views of the National Science Foundation. We also acknowledge support from Army Research Office contract # W911NF-23-1-0327.

## Source Code and Data

Code and videos for the analyses described herein is publicly available at: https://github.com/caitlingrasso/bio-connectivity.git

## References

1. Larsen AZ, Kummer U. In: Information Processing in Calcium Signal Transduction. Springer Berlin Heidelberg; 2003. p. 153–178.

2. Kudla J, BatističO, Hashimoto K. Calcium Signals: The Lead Currency of Plant Information Processing. The Plant Cell. 2010;22(3):541–563. doi:10.1105/tpc.109.072686.

3. Ingber DE. Tensegrity II. How structural networks influence cellular information processing networks. Journal of Cell Science. 2003;116(8):1397–1408. doi:10.1242/jcs.00360.

4. Balázsi G, van Oudenaarden A, Collins J. Cellular Decision Making and Biological Noise: From Microbes to Mammals. Cell. 2011;144(6):910–925. doi:10.1016/j.cell.2011.01.030.

5. Pezzulo G, D’Amato L, Mannella F, Priorelli M, Van de Maele T, Stoianov IP, et al. Neural representation in active inference: using generative models to interact with – and understand – the lived world; 2023.

6. Daniels BC, Flack JC, Krakauer DC. Dual Coding Theory Explains Biphasic Collective Computation in Neural Decision-Making. Frontiers in Neuroscience. 2017;11. doi:10.3389/fnins.2017.00313.

7. Schneidman E, Bialek W, Berry MJ. An information theoretic approach to the functional classification of neurons. 2002; doi:10.48550/ARXIV.PHYSICS/0212114.

8. Palmer SE, Marre O, Berry MJ, Bialek W. Predictive information in a sensory population. Proceedings of the National Academy of Sciences. 2015;112(22):6908–6913. doi:10.1073/pnas.1506855112.

9. Li M, Han Y, Aburn MJ, Breakspear M, Poldrack RA, Shine JM, et al. Transitions in information processing dynamics at the whole-brain network level are driven by alterations in neural gain. PLOS Computational Biology. 2019;15(10):e1006957. doi:10.1371/journal.pcbi.1006957.

10. Lyon P, Keijzer F, Arendt D, Levin M. Reframing cognition: getting down to biological basics. Philosophical Transactions of the Royal Society B: Biological Sciences. 2021;376(1820):20190750. doi:10.1098/rstb.2019.0750.

11. Baluška F, Levin M. On Having No Head: Cognition throughout Biological Systems. Frontiers in Psychology. 2016;7. doi:10.3389/fpsyg.2016.00902.

12. Moore D, Walker SI, Levin M. Cancer as a disorder of patterning information: computational and biophysical perspectives on the cancer problem. Convergent Science Physical Oncology. 2017;3(4):043001. doi:10.1088/2057-1739/aa8548.

13. Hoel E, Levin M. Emergence of informative higher scales in biological systems: a computational toolkit for optimal prediction and control. Communicative and Integrative Biology. 2020;13(1):108–118. doi:10.1080/19420889.2020.1802914.

14. Kudithipudi D, Aguilar-Simon M, Babb J, Bazhenov M, Blackiston D, Bongard J, et al. Biological underpinnings for lifelong learning machines. Nature Machine Intelligence. 2022;4(3):196–210. doi:10.1038/s42256-022-00452-0.

15. Ghosheh M, Ehrlich A, Ioannidis K, Ayyash M, Goldfracht I, Cohen M, et al. Electro-metabolic coupling in multi-chambered vascularized human cardiac organoids. Nature Biomedical Engineering. 2023;7(11):1493–1513. doi:10.1038/s41551-023-01071-9.

16. Spira ME, Hai A. Multi-electrode array technologies for neuroscience and cardiology. Nature Nanotechnology. 2013;8(2):83–94. doi:10.1038/nnano.2012.265.

17. Simons M, Mlodzik M. Planar Cell Polarity Signaling: From Fly Development to Human Disease. Annual Review of Genetics. 2008;42(1):517–540. doi:10.1146/annurev.genet.42.110807.091432.

18. Butler MT, Wallingford JB. Planar cell polarity in development and disease. Nature Reviews Molecular Cell Biology. 2017;18(6):375–388. doi:10.1038/nrm.2017.11.

19. Rong G, Corrie SR, Clark HA. In Vivo Biosensing: Progress and Perspectives. ACS Sensors. 2017;2(3):327–338. doi:10.1021/acssensors.6b00834.

20. Jones C, Roper VC, Foucher I, Qian D, Banizs B, Petit C, et al. Ciliary proteins link basal body polarization to planar cell polarity regulation. Nature Genetics. 2007;40(1):69–77. doi:10.1038/ng.2007.54.

21. Dos Santos M, Backer S, Saintpierre B, Izac B, Andrieu M, Letourneur F, et al. Single-nucleus RNA-seq and FISH identify coordinated transcriptional activity in mammalian myofibers. Nature Communications. 2020;11(1). doi:10.1038/s41467-020-18789-8.

22. Petrany MJ, Swoboda CO, Sun C, Chetal K, Chen X, Weirauch MT, et al. Single-nucleus RNA-seq identifies transcriptional heterogeneity in multinucleated skeletal myofibers. Nature Communications. 2020;11(1). doi:10.1038/s41467-020-20063-w.

23. Jin S, Guerrero-Juarez CF, Zhang L, Chang I, Ramos R, Kuan CH, et al. Inference and analysis of cell-cell communication using CellChat. Nature Communications. 2021;12(1). doi:10.1038/s41467-021-21246-9.

24. Rubinov M, Sporns O. Complex network measures of brain connectivity: Uses and interpretations. NeuroImage. 2010;52(3):1059–1069. doi:10.1016/j.neuroimage.2009.10.003.

25. Friston KJ. Functional and effective connectivity: a review. Brain Connectivity. 2011;1(1):13–36. doi:10.1089/brain.2011.0008.

26. Shannon CE. A Mathematical Theory of Communication. Bell System Technical Journal. 1948;27(3):379–423. doi:10.1002/j.1538-7305.1948.tb01338.x.

27. Schreiber T. Measuring Information Transfer. Physical Review Letters. 2000;85(2):461–464. doi:10.1103/PhysRevLett.85.461.

28. Brodskiy PA, Zartman JJ. Calcium as a signal integrator in developing epithelial tissues. Physical Biology. 2018;15(5):051001. doi:10.1088/1478-3975/aabb18.

29. Dodd AN, Kudla J, Sanders D. The Language of Calcium Signaling. Annual Review of Plant Biology. 2010;61(1):593–620. doi:10.1146/annurev-arplant-070109-104628.

30. McMillen P, Walker SI, Levin M. Information Theory as an Experimental Tool for Integrating Disparate Biophysical Signaling Modules. International Journal of Molecular Sciences. 2022;23(17). doi:10.3390/ijms23179580.

31. Lansdown ABG. Calcium: a potential central regulator in wound healing in the skin. Wound Repair and Regeneration. 2002;10(5):271–285. doi:10.1046/j.1524-475x.2002.10502.x.

32. Knot HJ. Twenty Years of Calcium Imaging: Cell Physiology to Dye For. Molecular Interventions. 2005;5(2):112–127. doi:10.1124/mi.5.2.8.

33. Li S, Liu Y, Zhang N, Li W, Xu Wj, Xu Yq, et al. Perspective of Calcium Imaging Technology Applied to Acupuncture Research. Chinese Journal of Integrative Medicine. 2023;30(1):3–9. doi:10.1007/s11655-023-3692-2.

34. Subramaniam T, Fauzi MB, Lokanathan Y, Law JX. The Role of Calcium in Wound Healing. International Journal of Molecular Sciences. 2021;22(12):6486. doi:10.3390/ijms22126486.

35. Aihara E, Hentz CL, Korman AM, Perry NPJ, Prasad V, Shull GE, et al. In Vivo Epithelial Wound Repair Requires Mobilization of Endogenous Intracellular and Extracellular Calcium. Journal of Biological Chemistry. 2013;288(47):33585–33597. doi:10.1074/jbc.m113.488098.

36. Zulueta-Coarasa T, Fernandez-Gonzalez R. Tension (re)builds: Biophysical mechanisms of embryonic wound repair. Mechanisms of Development. 2017;144:43–52. doi:10.1016/j.mod.2016.11.004.

37. Yoo SK, Freisinger CM, LeBert DC, Huttenlocher A. Early redox, Src family kinase, and calcium signaling integrate wound responses and tissue regeneration in zebrafish. Journal of Cell Biology. 2012;199(2):225–234. doi:10.1083/jcb.201203154.

38. Jones EA, Woodland HR. Development of the ectoderm in Xenopus: Tissue specification and the role of cell association and division. Cell. 1986;44(2):345–355. doi:10.1016/0092-8674(86)90769-5.

39. Green J. In: The Animal Cap Assay. Humana Press;. p. 1–14.

40. Lee J, Møller AF, Chae S, Bussek A, Park TJ, Kim Y, et al. A single-cell, time-resolved profiling of Xenopus mucociliary epithelium reveals nonhierarchical model of development. Science Advances. 2023;9(14). doi:10.1126/sciadv.add5745.

41. Walentek P, Bogusch S, Thumberger T, Vick P, Dubaissi E, Beyer T, et al. A novel serotonin-secreting cell type regulates ciliary motility in the mucociliary epidermis of Xenopus tadpoles. Development. 2014;141(7):1526–1533. doi:10.1242/dev.102343.

42. Walentek P, Quigley IK. What we can learn from a tadpole about ciliopathies and airway diseases: Using systems biology in Xenopus to study cilia and mucociliary epithelia. genesis. 2017;55(1–2). doi:10.1002/dvg.23001.

43. Walentek P. Xenopus epidermal and endodermal epithelia as models for mucociliary epithelial evolution, disease, and metaplasia. genesis. 2021;59(1–2). doi:10.1002/dvg.23406.

44. Dubaissi E, Papalopulu N. Embryonic frog epidermis: a model for the study of cell-cell interactions in the development of mucociliary disease. Disease Models and Mechanisms. 2011;4(2):179–192. doi:10.1242/dmm.006494.

45. Davidson LA, Ezin AM, Keller R. Embryonic wound healing by apical contraction and ingression in Xenopus laevis. Cell Motility. 2002;53(3):163–176. doi:10.1002/cm.10070.

46. S Kriegman MLJB D Blackiston. A Scalable Pipeline for Designing Reconfigurable Organisms. Proc Natl Acad Sci U S A. 2020;117(4):1853–1859. doi:10.1073/pnas.1910837117.

47. Blackiston D, Kriegman S, Bongard J, Levin M. Biological Robots: Perspectives on an Emerging Interdisciplinary Field. Soft Robotics. 2023;10(4):674–686. doi:10.1089/soro.2022.0142.

48. Werner ME, Mitchell BJ. In: Using Xenopus Skin to Study Cilia Development and Function. Elsevier; 2013. p. 191–217.

49. Mitchell B, Stubbs JL, Huisman F, Taborek P, Yu C, Kintner C. The PCP Pathway Instructs the Planar Orientation of Ciliated Cells in the Xenopus Larval Skin. Current Biology. 2009;19(11):924–929. doi:10.1016/j.cub.2009.04.018.

50. Deblandre G, Wettstein DA, Koyano-Nakagawa N, Kintner C. A two-step mechanism generates the spacing pattern of the ciliated cells in the skin of Xenopus embryos. Development. 1999;126(21):4715–4728. doi:10.1242/dev.126.21.4715.

51. Collins C, Ventrella R, Mitchell BJ. In: Building a ciliated epithelium: Transcriptional regulation and radial intercalation of multiciliated cells. Elsevier; 2021. p. 3–39.

52. Walentek P. In: Manipulating and Analyzing Cell Type Composition of the Xenopus Mucociliary Epidermis. Springer New York; 2018. p. 251–263.

53. Blackiston D, Lederer E, Kriegman S, Garnier S, Bongard J, Levin M. A Cellular Platform for the Development of Synthetic Living Machines. Science Robotics. 2021;6(52):eabf1571. doi:10.1126/scirobotics.abf1571.

54. Avants B, Tustison NJ, Song G. Advanced Normalization Tools: V1.0. The Insight Journal. 2009; doi:10.54294/uvnhin.

55. Stringer C, Wang T, Michaelos M, Pachitariu M. Cellpose: a generalist algorithm for cellular segmentation. Nature Methods. 2020;18(1):100–106. doi:10.1038/s41592-020-01018-x.

56. Daube C, Gross J, Ince RAA. A whitening approach for Transfer Entropy permits the application to narrow-band signals. arXiv: 220102461 [q-bio]. 2022;.

57. Liu TT, Nalci A, Falahpour M. The global signal in fMRI: Nuisance or Information? NeuroImage. 2017;150:213–229. doi:10.1016/j.neuroimage.2017.02.036.

58. Zamani Esfahlani F, Jo Y, Faskowitz J, Byrge L, Kennedy DP, Sporns O, et al. High-amplitude cofluctuations in cortical activity drive functional connectivity. Proceedings of the National Academy of Sciences. 2020;117(45):28393–28401. doi:10.1073/pnas.2005531117.

59. Pope M, Fukushima M, Betzel RF, Sporns O. Modular origins of high-amplitude cofluctuations in fine-scale functional connectivity dynamics. Proceedings of the National Academy of Sciences. 2021;118(46). doi:10.1073/pnas.2109380118.

60. Sapir L, Tzlil S. Talking over the extracellular matrix: How do cells communicate mechanically? Seminars in Cell and Developmental Biology. 2017;71:99–105. doi:10.1016/j.semcdb.2017.06.010.

61. Hervé JC, Derangeon M. Gap-junction-mediated cell-to-cell communication. Cell and Tissue Research. 2012;352(1):21–31. doi:10.1007/s00441-012-1485-6.

62. Jeub LGS, Sporns O, Fortunato S. Multiresolution Consensus Clustering in Networks. Scientific Reports. 2018;8(1). doi:10.1038/s41598-018-21352-7.

63. Blondel VD, Guillaume JL, Lambiotte R, Lefebvre E. Fast unfolding of communities in large networks. J Stat Mech (2008) P10008. 2008;2008(10):P10008. doi:10.1088/1742-5468/2008/10/p10008.

64. Rubinov M. Circular and unified analysis in network neuroscience. eLife. 2023;12. doi:10.7554/elife.79559.

65. Power J, Schlaggar B, Lessov-Schlaggar C, Petersen S. Evidence for Hubs in Human Functional Brain Networks. Neuron. 2013;79(4):798–813. doi:10.1016/j.neuron.2013.07.035.

66. Rosas FE, Mediano PAM, Luppi AI, Varley TF, Lizier JT, Stramaglia S, et al. Disentangling high-order mechanisms and high-order behaviours in complex systems. Nature Physics. 2022;18(5):476–477. doi:10.1038/s41567-022-01548-5.

67. Battiston F, Cencetti G, Iacopini I, Latora V, Lucas M, Patania A, et al. Networks beyond pairwise interactions: Structure and dynamics. Physics Reports. 2020;874:1–92. doi:10.1016/j.physrep.2020.05.004.

68. Newman EL, Varley TF, Parakkattu VK, Sherrill SP, Beggs JM. Revealing the Dynamics of Neural Information Processing with Multivariate Information Decomposition. Entropy. 2022;24(7):930. doi:10.3390/e24070930.

69. Varley TF, Pope M, Grazia M, Joshua, Sporns O. Partial entropy decomposition reveals higher-order information structures in human brain activity. Proceedings of the National Academy of Sciences. 2023;120(30). doi:10.1073/pnas.2300888120.

70. Varley TF, Pope M, Faskowitz J, Sporns O. Multivariate information theory uncovers synergistic subsystems of the human cerebral cortex. Communications Biology. 2023;6(1). doi:10.1038/s42003-023-04843-w.

71. Brugués A, Anon E, Conte V, Veldhuis JH, Gupta M, Colombelli J, et al. Forces driving epithelial wound healing. Nature Physics. 2014;10(9):683–690. doi:10.1038/nphys3040.

72. Faber J. Normal Table of Xenopus Laevis (Daudin). Nieuwkoop PD, editor. Milton: CRC Press LLC; 1994.

73. Chen J, Xia L, Bruchas MR, Solnica-Krezel L. Imaging early embryonic calcium activity with GCaMP6s transgenic zebrafish. Developmental Biology. 2017;430(2):385–396. doi:10.1016/j.ydbio.2017.03.010.

74. Offner T, Daume D, Weiss L, Hassenklöver T, Manzini I. Whole-Brain Calcium Imaging in Larval Xenopus. Cold Spring Harbor Protocols. 2020;2020(12):pdb.prot106815. doi:10.1101/pdb.prot106815.

75. Werner ME, Mitchell BJ. Understanding ciliated epithelia: The power of Xenopus. genesis. 2011;50(3):176–185. doi:10.1002/dvg.20824.

76. Spassky N, Meunier A. The development and functions of multiciliated epithelia. Nature Reviews Molecular Cell Biology. 2017;18(7):423–436. doi:10.1038/nrm.2017.21.

77. Huynh MH, Hong H, Delovitch S, Desser S, Ringuette M. Association of SPARC (osteonectin, BM-40) with extracellular and intracellular components of the ciliated surface ectoderm of Xenopus embryos. Cell motility and the cytoskeleton. 2000;47:154–162. doi:10.1002/1097-0169(200010)47:2¡154::AID-CM6¿3.0.CO;2-L.

78. Angerilli A, Smialowski P, Rupp R. The Xenopus animal cap transcriptome: building a mucociliary epithelium. Nucleic Acids Research. 2018;46(17):8772–8787. doi:10.1093/nar/gky771.

79. Kang HJ, Kim HY. Mucociliary Epithelial Organoids from Xenopus Embryonic Cells: Generation, Culture and High-Resolution Live Imaging. Journal of Visualized Experiments. 2020;(161). doi:10.3791/61604.

80. Sive HL, Grainger RM, Harland RM. Animal Cap Isolation from Xenopus laevis:Figure 1. Cold Spring Harbor Protocols. 2007;2007(6):pdb.prot4744. doi:10.1101/pdb.prot4744.

81. Schindelin J, Arganda-Carreras I, Frise E, Kaynig V, Longair M, Pietzsch T, et al. Fiji: an open-source platform for biological-image analysis. Nature Methods. 2012;9(7):676–682. doi:10.1038/nmeth.2019.

82. Bossomaier T, Barnett L, Harré M, Lizier JT. An Introduction to Transfer Entropy: Information Flow in Complex Systems. Springer; 2016.

83. Weiser SC, Mullen BR, Ascencio D, Ackman JB. Data-driven segmentation of cortical calcium dynamics. PLOS Computational Biology. 2023;19(5):e1011085. doi:10.1371/journal.pcbi.1011085.

84. Murphy MC, Chan KC, Kim SG, Vazquez AL. Macroscale variation in resting-state neuronal activity and connectivity assessed by simultaneous calcium imaging, hemodynamic imaging and electrophysiology. NeuroImage. 2018;169:352–362. doi:10.1016/j.neuroimage.2017.12.070.

85. Afyouni S, Smith SM, Nichols TE. Effective degrees of freedom of the Pearson’s correlation coefficient under autocorrelation. NeuroImage. 2019;199:609–625. doi:10.1016/j.neuroimage.2019.05.011.

86. Lizier JT. The Local Information Dynamics of Distributed Computation in Complex Systems. Springer Theses. Berlin, Heidelberg: Springer Berlin Heidelberg; 2013. Available from: http://link.springer.com/10.1007/978-3-642-32952-4.

87. Wibral M, Lizier J, Vögler S, Priesemann V, Galuske R. Local active information storage as a tool to understand distributed neural information processing. Frontiers in Neuroinformatics. 2014;8. doi:10.3389/fninf.2014.00001.

88. Lizier JT. JIDT: An Information-Theoretic Toolkit for Studying the Dynamics of Complex Systems. Frontiers in Robotics and AI. 2014;1. doi:10.3389/frobt.2014.00011.

89. Wollstadt P, Lizier JT, Vicente R, Finn C, Martinez-Zarzuela M, Mediano P, et al. IDTxl: The Information Dynamics Toolkit xl: a Python package for the efficient analysis of multivariate information dynamics in networks. Journal of Open Source Software. 2019;4(34):1081. doi:10.21105/joss.01081.

